# Heat inactivation of the Monkeypoxvirus

**DOI:** 10.1101/2022.08.10.502482

**Authors:** Christophe Batéjat, Quentin Grassin, Maxence Feher, Damien Hoinard, Jessica Vanhomwegen, Jean-Claude Manuguerra, India Leclercq

## Abstract

Different kinds of media spiked with Monkeypoxvirus (MPXV) were subjected to heat inactivation at different temperatures, for various periods of time. Our results showed that MPXV was inactivated in less than 5 min at 70 °C and in less than 15 min at 60 °C, with no difference between virus from West African and Central African clades. Such indications could help laboratory workers to manipulate the virus in optimal biosafety conditions and improve their protocols.

## Introduction

Since April 2022, thousands of monkeypox cases have been reported in non-endemic countries. This unprecedented outbreak is caused by a monkeypoxvirus (MPXV) belonging to the *Poxviridae* family and *Orthopoxvirus* genus. The majority of confirmed cases (n=31,800 the 9^th^ of August 2022) are from the WHO European Region (1). Men who have sex with men who have reported recent sex with new and multiple partners are predominantly infected. Since the discovery of the virus in 1958, growing numbers of monkeypox cases were reported over the past five decades, in 10 African countries and 4 countries outside of Africa. Observing such a huge number of cases outside Africa is more than unusual and reinforce the necessity to better characterize this virus and its transmission and pathogenesis. Manipulation of clinical samples and research on the virus are necessary to help fighting and stop the current outbreak. Viral inactivation procedures are therefore necessary to allow safe experimental laboratory conditions. Heat treatment is a widely used inactivation method for viruses. Personal Protective Equipment (PPE), laboratory materials, hospital equipment, transportation media, and biological samples are often heated to inactivate viruses. Heat is thought to denature secondary structures of various molecules constituting the virus. Data on orthopoxviruses are only available for vaccinia virus and variola virus, but not for MPXV. The necessary time for achieving at least a 4 log10 reduction in citrate-phosphate buffer was 15 min at 60°C and 90 min at 50°C (2). Due to their close structure, Vaccinia virus and MPXV might have a similar susceptibility towards thermal inactivation, as previously observed with biocidal agent (3). To address this question, we submitted two different strains of MPXV to temperatures commonly used in laboratory for viral inactivation, during various periods of time and tested them for residual infectivity by plaque assay method.

## Materials and Methods

### Cell lines and viruses

African green monkey cells (Vero E6) were grown in Dulbecco’s modified Eagle’s medium (DMEM 1X, GIBCO) supplemented with 5% fetal calf serum (FCS) and antibiotics (0.1 unit penicillin, 0.1 mg.mL^-1^ streptomycin, GIBCO) at 37 °C in a humidified 5% CO_2_ incubator.

A strain of MPXV isolated in the Congo basin (named LK for Lokole region) was also grown on Vero E6 cells for 3 passages with a titer of 3.49×10^7^ PFU.mL^-1^ (Plaque Forming Unit.mL^-1^). A human strain of MPXV, isolated from a French patient in June 2022 (strain MPXV/2022/FR/CMIP named CMIP2022), was grown on Vero E6 cells for 2 passages. The virus was titrated by plaque assay and the titer was 3.4×10^7^ PFU.mL^-1^.

### Heat inactivation

MPXV strains were diluted ½ in two different kinds of media that is Viral Transport Media (VTM; 330C, Copan) and FCS (F2442, Sigma-Aldrich). Two hundreds microliters of each sample were submitted in triplicate to various temperatures for different periods of time, ranging from 30 sec to 90 min. Samples were inactivated in a calibrated and verified dry water bath, cooled on ice and tested for infectivity by using a plaque assay procedure adapted from Matrosovich et al. (4). The temperatures tested were 56°C, 60°C, 70°C and 95°C to mimic commonly temperature used for serosurveys, inactivation of high treat pathogens, standard temperature used to inactivate the MPXV prior for diagnosis, and temperature used for rapid inactivation of viruses and compatible with molecular biology techniques, respectively.

All experiments were conducted under strict BSL3 conditions.

### Plaque assay

Vero E6 cells were grown in 6-well plates to reach the appropriate confluency. Cells were washed once with Dulbecco’s Phosphate-Buffered Saline (DPBS) then infected with 10-fold dilution of heat inactivated samples. After 1 hour and a half of adsorption at 37°C, 3 mL of DMEM containing 1.2% Avicel (RC581; FMC BioPolymer) were added to the cells. Plaques were revealed after 5 days of incubation at 37°C by staining with crystal violet and paraformaldehyde.

### Quantitative PCR

Viral DNA of samples with the longest exposure time were extracted using NucleoSpin^®^ 96 Virus Core kit following the manufacturer’s instructions. Viral DNA was amplified using SsoAdvanced Universal Probes Supermix (BioRad) kit with the following generic primers and probe targeting the G2R gene (5): Forward Primer: 5’- GGAAAATGTAAAGACAACGAATACAG – 3’;Reverse Primer: 5’- GCTATCACATAATCTGGAAGCGTA –3’; Hydrolysis probe: 5’- AAGCCGTAATCTATGTTGTCTATCGTGTCC – 3’. Cycle thresholds (Ct) obtained for each condition were compared to the initial Ct obtained with the virus suspension before its heat treatment.

## Results and discussion

VTM and FCS were spiked with two different strains of MPXV, one belonging to the West African clade (CMIP2022) and the other to the Central African clade (LK) both with titers around 1×10^7^ PFU.mL^-1^. All samples were submitted in triplicate to different temperatures for various periods of time in a calibrated dry water bath. The results are summarized in Tables 1 and 2. They showed that both strains of MPXV were inactivated in all cases, except at 95°C after 30 s of heat treatment and after 30 min at 56°C. Thirty seconds at 95°C was probably not long enough to reach such temperature inside the tube, as already observed with Syndrome Acute Respiratory Virus 2 (SARS-CoV-2) (6). As reaching of targeted temperature could take more time with higher volumes than 200 μL, we recommend to treat samples during 30 min at 60°C to inactivate the virus, although heat treatment during 15 min at 60°C of 500 μL of non-diluted viral suspensions with titers of 5.25×10^7^ (CMIP2022) and 3.49×10^7^ PFU.mL^-1^ (LK) showed complete viral inactivation (data not shown). Recommended times and temperature of 70°C in some extraction kits are also efficient to allow complete virus inactivation. Our data are in agreement with previous results obtained with vaccinia virus at 60°C (7). However, MPXV was still infectious after 30 min at 56°C, suggesting that thermal inactivation of serum products must be performed for at least one hour at this temperature to reach full inactivation of high titer sera.

**Table 1:**
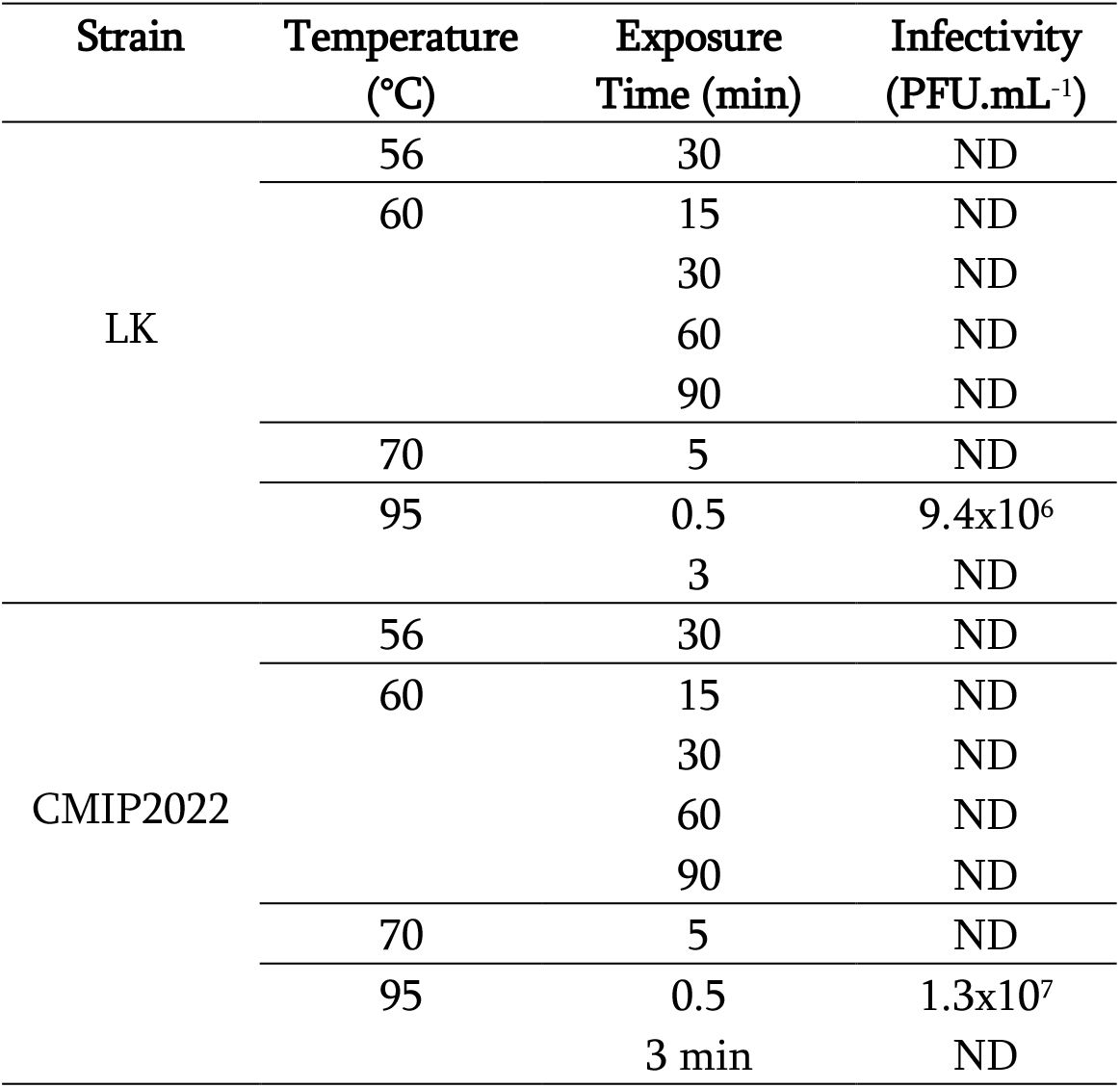
Viral titers obtained after heat inactivation of viruses diluted in FCS. ND: not detected (below the limit of virus detection, which corresponded to 25 PFU.mL^-1^).

**Table 2:**
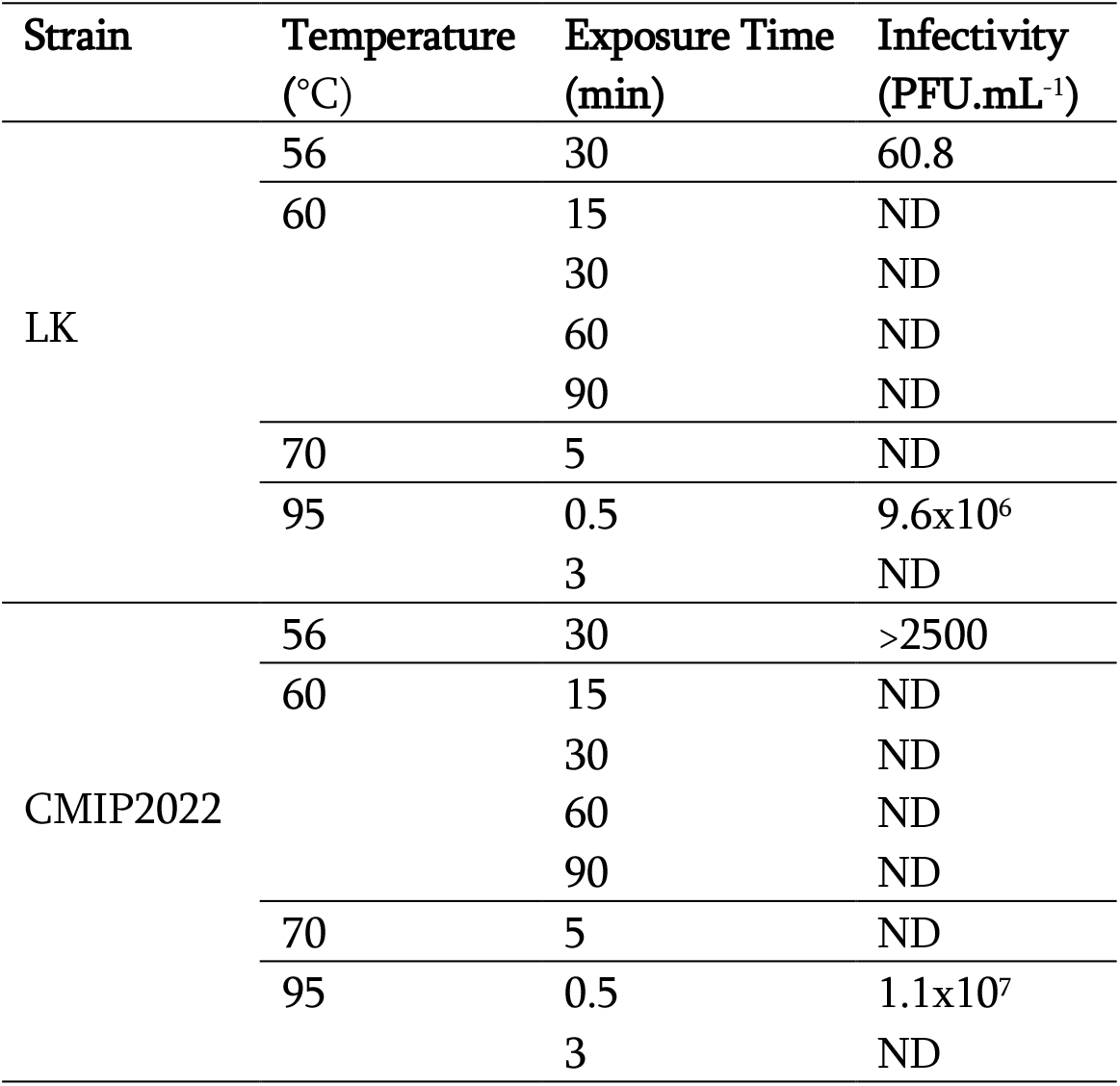
Viral titers obtained after heat inactivation of viruses diluted in VTM. ND: not detected (below the limit of virus detection, which corresponded to 25 PFU.mL^-1^).

Quantitative PCR (qPCR) was performed in order to explore the impact of heat treatment on viral genome detection. Results summarized in table 3 showed that heat treatment has a very weak impact on subsequent viral DNA detection by qPCR, with variations of Ct not exceeding 2.09 at 60°C and 70°C, and 3.46 at 95°C, suggesting that the majority of viral DNA remained intact in virus particles.

**Table 3:**
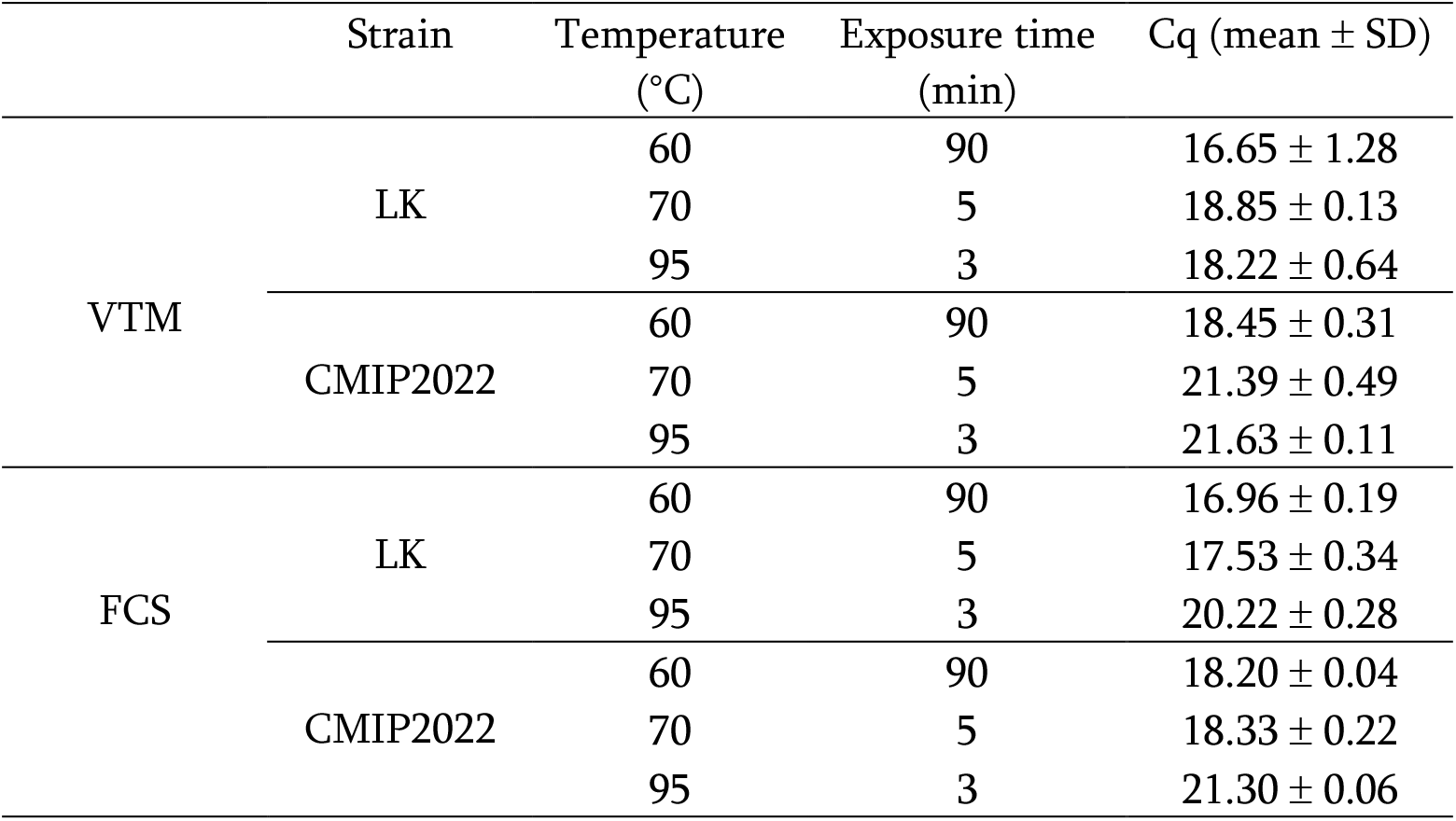
Quantification of viral DNA in heat inactivated samples. Mean cycle quantification (Cq) values were obtained for each of the longest inactivated condition done in triplicate (± Standard deviation). Initial Cq were 16.76 ±.11 and 20.19 ±.18 for LK and CMIP2022 strains respectively.

MPXV is thus relatively sensitive to heat inactivation under laboratory conditions and such results can help workers to improve their protocols, and is a start for knowledge and comprehension of MPXV survival mechanisms outside the host. The data described in this paper were obtained with viruses in suspension in VTM and FCS, and submitted to high temperatures and are useful for laboratory protocols. Further experimental data are necessary to evaluate the persistence of MPXV on various matrices at different temperatures and evaluate the potential role of contaminated surfaces in the virus transmission.

## CRediT authorship contribution statement

Christophe Batéjat: Conceptualization, Formal analysis, Investigation, Methodology, Project administration, Resources, Supervision, Validation, Writing - review & editing. Quentin Grassin: Investigation. Maxence Feher: Investigation. Damien Hoinard: Investigation. Jessica Vanhomwegen: Writing - review & editing. Jean-Claude Manuguerra: Resources, Writing - review & editing. India Leclercq: Conceptualization, Formal analysis, Investigation, Methodology, Project administration, Resources, Supervision, Validation, Writing - original draft, Writing - review & editing.

The Authors declare that there is no conflict of interest.

## Notes

### Competing Interest Statement

The authors have declared no competing interest.

