## Supplementary material for "Heat inactivation of the Monkeypoxvirus": Tables

**Table 1:** Viral titers obtained after heat inactivation of viruses diluted in FCS. ND: not detected (below the limit of virus detection, which corresponded to 25 PFU.mL$^{-1}$).

| Strain | Temperature (°C) | Exposure Time (min) | Infectivity (PFU.mL$^{-1}$) |
|---|---|---|---|
| LK | 56 | 30 | ND |
| | 60 | 15 | ND |
| | | 30 | ND |
| | | 60 | ND |
| | | 90 | ND |
| | 70 | 5 | ND |
| | 95 | 0.5 | $9.4x10^6$ |
| | | 3 | ND |
| CMIP2022 | 56 | 30 | ND |
| | 60 | 15 | ND |
| | | 30 | ND |
| | | 60 | ND |
| | | 90 | ND |
| | 70 | 5 | ND |
| | 95 | 0.5 | $1.3x10^7$ |
| | | 3 min | ND |

**Table 2:** Viral titers obtained after heat inactivation of viruses diluted in VTM. ND: not detected (below the limit of virus detection, which corresponded to 25 PFU.mL$^{-1}$).

| Strain | Temperature (°C) | Exposure Time (min) | Infectivity (PFU.mL$^{-1}$) |
|---|---|---|---|
| LK | 56 | 30 | 60.8 |
| | 60 | 15 | ND |
| | | 30 | ND |
| | | 60 | ND |
| | | 90 | ND |
| | 70 | 5 | ND |
| | 95 | 0.5 | $9.6x10^6$ |
| | | 3 | ND |
| CMIP2022 | 56 | 30 | >2500 |
| | 60 | 15 | ND |
| | | 30 | ND |
| | | 60 | ND |
| | | 90 | ND |
| | 70 | 5 | ND |
| | 95 | 0.5 | $1.1x10^7$ |
| | | 3 | ND |

**Table 3:** Quantification of viral DNA in heat inactivated samples. Mean cycle quantification (Cq) values were obtained for each of the longest inactivated condition done in triplicate (± Standard deviation). Initial Cq were 16.76 ± .11 and 20.19 ± .18 for LK and CMIP2022 strains respectively.

| | Strain | Temperature (°C) | Exposure time (min) | Cq (mean ± SD) |
|---|---|---|---|---|
| VTM | LK | 60 | 90 | 16.65 ± 1.28 |
| | | 70 | 5 | 18.85 ± 0.13 |
| | | 95 | 3 | 18.22 ± 0.64 |
| | CMIP2022 | 60 | 90 | 18.45 ± 0.31 |
| | | 70 | 5 | 21.39 ± 0.49 |
| | | 95 | 3 | 21.63 ± 0.11 |
| FCS | LK | 60 | 90 | 16.96 ± 0.19 |
| | | 70 | 5 | 17.53 ± 0.34 |
| | | 95 | 3 | 20.22 ± 0.28 |
| | CMIP2022 | 60 | 90 | 18.20 ± 0.04 |
| | | 70 | 5 | 18.33 ± 0.22 |
| | | 95 | 3 | 21.30 ± 0.06 |
